# Rooting morphologically divergent taxa – slow-evolving sequence data might help

**DOI:** 10.1101/2020.03.15.983684

**Authors:** Jorge Flores, Alexander C. Bippus, Alexandru Tomescu, Neil Bell, Jaakko Hyvönen

## Abstract

When fossils are sparse and the lineages studied are very divergent morphologically, analyses based exclusively on morphology may lead to conflicting and unexpected hypotheses. Through integration of data from conservative genes/gene regions the terminals including these data can anchor or constrain the search, thereby practically circumscribing the search space of the combined analyses. In this study, we revisit the phylogeny of a highly divergent group of mosses, class Polytrichopsida. We supplemented the morphological matrix by adding sequence data of the nuclear gene 18S, chloroplast genes *rbc*L and *rps*4, plus the mitochondrial gene *nad*5. For the phylogenetic analyses we used parsimony as the optimality criterion. Analyses that included all the terminals resulted in one most parsimonious tree with a clade comprised of *Alophosia azorica* and the fossil *Meantoinea alophosioides* representing the basal-most lineage. Analyses with different outgroup sampling produced the same topology for most ingroup relationships. An analysis excluding morphological characters and the four terminals for which only morphological characters were scored (the two fossil and two extant terminals) resulted in one optimal tree with identical topology to the one obtained when including all terminals. These results are largely congruent with those obtained in the recent analyses based exclusively on sequence level data of a larger number of terminals. Our results indicate that large size and complexity of the gametophyte have evolved independently in several lineages. Notably, the nodes of the backbone of the most parsimonious tree have very low support values, thus these inferred relationships could change if new additional information conflicts with the current data. Future studies should be aimed at incorporating all terminals into phylogenetic analyses, which is not an unrealistic goal for a group with less than 200 species. Also, additional fossils, some of which await detailed examination and description, need to be included. Whether these will affect the overall pattern of phylogeny presented here remains to be seen. In a group that is obviously very ancient, we cannot assume, *a priori*, that currently known fossil taxa, which go back in time less than 140 Ma, represent the oldest lineages of the group.

---

Our understanding of organismal diversity and the most robust hypotheses about its history suggest that evolution by default produces divergence. It is also clear that extant organisms represent only a small fraction of the total biodiversity that has lived on this planet throughout the more than 3.5 Ga history of life (e.g. Marshall, 2017), thus, extinction was a key factor shaping extant diversity. Such rampant extinction leads to situations where defining homologies can be difficult. In turn, this leads to difficulties in phylogenetic analyses. Fossils are one the most widely recognized sources of information that can provide more insight in these situations. Numerous classic cases highlighting the importance of the fossils are provided, for example, by Forey & al. (1992), Smith (1994), Crane & al. (2004), Hilton & Bateman (2006), and Rothwell & Nixon (2006). On the other hand, when fossils are sparse and the lineages studied are very divergent morphologically, hypotheses based exclusively on morphology may lead to conflicting and unexpected hypotheses (e.g. Bippus & al., 2018). Sequence data of slowly evolving regions of the genome could offer a remedy in such cases. It is widely recognized that rates of evolution do vary between characters and, thus, when morphology is exceptionally divergent between lineages, at least some gene regions may still be highly similar. We know, for example, that the nuclear gene coding for 18S RNA is highly conserved, enabling comparisons of organisms even across major lineages (e.g. Hedderson & al., 1996, Hovmöller & al., 2002). Likewise, the chloroplast gene coding for the large subunit of rubisco (*rbc*L) is broadly conserved and was among the first genes widely used in phylogenetic studies of plants (e.g. Chase & al., 1993). The use of such highly conservative (i.e. slow-evolving) genes enables alignment of sequences and unequivocal assumptions about homology at the level of individual nucleotide positions. By using data from conservative genes/gene regions the terminals including these data can anchor or constrain the search, thereby practically circumscribing the search space of the combined analyses that integrate molecular and morphological data. As a result of this, more character state changes can be optimized on topologies; conversely, character state changes that were optimal based only on morphology might lose optimality upon combination of different data sources.

In this study, we revisit the phylogeny of a highly divergent group of mosses, class Polytrichopsida. We used a total evidence approach, following the example of Flores & al. (2017) wherein we supplemented the morphological matrix by adding sequence data into the analyses. Our analysis includes two fossils (*Eopolytrichum antiquum* Konopka & al. and *Meantoinea alophosioides* Bippus & al.) and over 40 extant species representing all genera of the Polytrichopsida, except for the recently described *Delongia* N. E. Bell & al. Because the class is morphologically isolated (i.e., highly divergent) from all other extant moss lineages, the root of the Polytrichopsida has been elusive, with different ingroup topologies recovered with different outgroup sampling regimes when only morphological characters are used in the phylogenetic analyses (Bippus & al., 2018). We now supplemented the morphological matrix with sequences of the nuclear gene 18S, chloroplast genes *rbc*L and *rps*4, plus the mitochondrial gene *nad*5. All these genes have been used in previous phylogenetic analyses of the group (e.g. Bell & Hyvönen, 2010, Hyvönen & al., 1998) and thus sequences for almost all the extant terminals included by Bippus & al. (2018) were readily available. Most of the sequences used were those used by Bell & Hyvönen (2010), with the exception of *rbc*L gene. Obtaining the whole sequence for this region was problematic and only about half of the gene sequence was available in Bell & Hyvönen (2010). Accordingly, we downloaded *rbc*L sequences from the GenBank as listed in the Appendix 1. Sequences of the genes listed above were also downloaded for the outgroup terminals. In cases where more than one sequence had been uploaded to the repository we used the longest sequence available. In order to avoid random attraction, or repulsion, of the terminals by sequences of unequal length that are due to sequencing artefacts, we were very conservative in our choice of the regions to be used. We excluded regions adjacent to the 5′and 3′ ends of the gene sequences where we had only few representatives. For *nad*5 we used only 492 nt from the 3′ end that also allowed alignment with the sequences obtained from the outgroup terminals. In order to maximize positional homology for the sequences downloaded we performed alignment using ClustalX (Larkin & al., 2007) under default settings. For the phylogenetic analyses we used parsimony as the optimality criterion. Discussions of strengths and weaknesses of different optimality criteria in phylogenetic analysis abound in the recent literature, and are beyond scope of this study. Nevertheless, we refer the reader to Flores & al. (2017), and particularly to Goloboff & al. (2018) for such discussions relative to analyses using morphological characters.

Initially, we removed parsimony uninformative characters from gene regions using the “mop uninformative characters” function of Winclada (Nixon, 2002). This produced a matrix of 519 molecular characters. Of these, 83 nt were from the 18S, 238 from the *rbc*L, 158 from the *rps*4, and 40 nt from the 3′ end of the mitochondrial *nad*5 gene. This dataset was combined with the matrix of Bippus & al. (2018), which includes 100 morphological characters, with 11 of which were coded as continuous and additive (Goloboff & al., 2006), while all the other (discrete) characters were treated as non-additive (unordered). The small number of terminals (45) enabled analysis using traditional search algorithms of the program TNT (Goloboff & Catalano 2016), with gaps treated as missing data. We performed analyses with different outgroup sampling regimes, similar to those of Bippus & al. (2018), i.e. with only *Alophosia azorica* Card. of Polytrichaceae to root the tree; or various samplings of the outgroup terminals, i.e. with using *Oedipodium griffithianum* Schwaegr.*, Sphagnum palustre* L. and *Tetraphis pellucida* Hedw, as outgroup terminals, or with the latter three supplemented by *Andreaea rupestris* Hedw., *Buxbaumia aphylla* Hedw., *Diphyscium foliosum* D. Mohr and *Funaria hygrometrica* Hedw. as outgroup terminals. All analyses were initiated with the random seed set to “0”, i.e. the CPU time is used as the random seed to randomize the order of the terminals. 1000 replicates of RAS (Wagner trees) were used in each search with 10 trees saved per replicate and TBR as a swapping algorithm. Tree searches were also performed with the same settings as used by Bippus & al. (2018). Support values were calculated using jackknife with the default values, i.e. with the results output as frequency differences (GC; Goloboff & al., 2003)

All analyses that included all the terminals included resulted in one optimal parsimonious tree with a length of 2089.52. In the other analyses we obtained exactly the same topology for most of the ingroup relationships. The only differences were present in the tree obtained using only three outgroup terminals (*Sphagnum palustre, Tetraphis pellucida* and *Oedipodium griffithianum*). In this tree, as compared to other trees, the fossil *Meantoinea alophosioides* was resolved as sister to the clade formed by two species of *Lyellia* R. Br., and not as sister to *Alophosia azorica*. Additionally, *Polytrichadelphus magellanicus* (Hedw.) Mitt. was resolved as sister to the clade formed by the two species of *Dawsonia* R. Br., while in the trees using other sampling sampling of the outgroup terminals *Polytrichadelphus* (Müll. Hal.) Mitt. and *Dawsonia* spp. formed a paraphyletic group basal to the large clade including most of the other genera (Fig. 1). An analysis excluding morphological characters and the four terminals for which only morphological characters were scored (the two fossil terminals plus *Pogonatum philippinense* (Broth.) Touw and *P. volvatum* (Müll. Hal.) Paris) resulted in one optimal tree with identical topology to the one obtained for all the 45 terminals.

**Figure 1.**
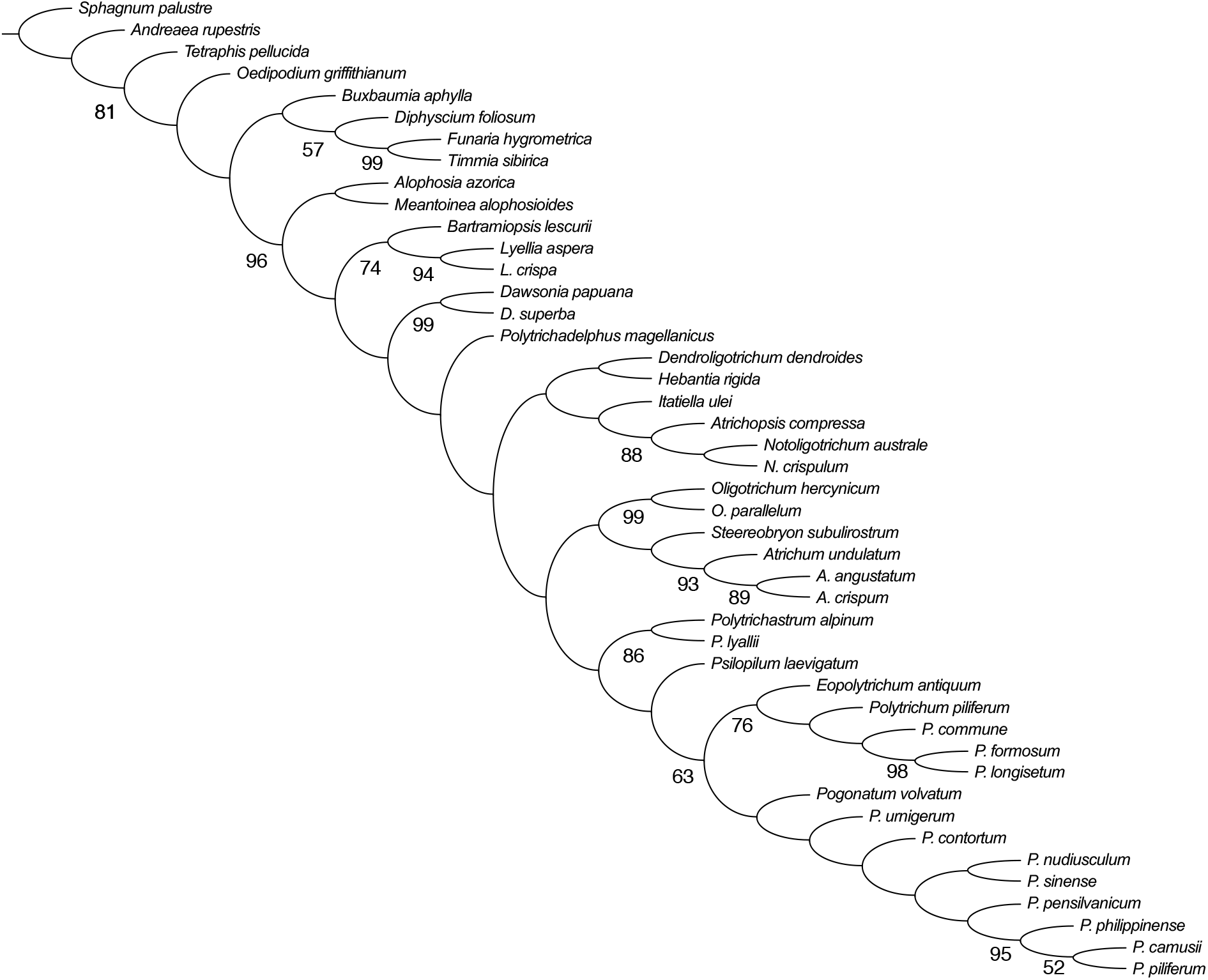
Tree obtained from combined analysis of the morphological (11 of these continuous) and sequence level (nc 18S 83, cp *rbc*L 238, *rps*4 158, mt *nad*5 40 nt) data, length 2089.52 steps. Jackknife support values >50 marked below branches.

It can be argued that in this analysis, morphology is swamped by sequence data because morphological characters are only ca. 16% of the total dataset. However, as discussed for example by Gatesy & al. (1999), the outcome of the total evidence analyses cannot be predicted from the sheer number, or proportions, of different types of characters and is, instead, determined by the amount of character congruence present in the dataset. Furthermore, as succinctly stated by Wheeler & al. (1993), the most logical explanation for the shared “signal” of diverse sources of information, whether morphological or sequence data, is their shared history. This perspective implies that combined analyses of molecular and morphological data provide the most robust hypotheses of relationships within a group (e.g. Flores & al., 2019). The results of our analyses show that the positions of the fossils and of the terminals that lack, at the moment, sequence level data are identical between the trees obtained with different outgroup sampling regimes. The only exception is the different placement of *Meantoinea alophosioides* and *Polytrichadelphus magellanicus* (Hedw.) Mitt. in a tree with only three outgroup terminals. However, it is important to note that this specific outgroup selection, which leaves out representatives of the largest clade of the peristomate mosses, cannot be defended. Therefore, we consider that the topology of Fig. 1 to represent our best estimate of the phylogeny of the major lineages of Polytrichopsida. This topology is largely congruent with the results obtained in analyses based exclusively on sequence level data of larger number of terminals (Bell & Hyvönen, 2010). When compared to the results of analyses based exclusively on morphological characters (Bippus & al., 2018), it is evident that the most obvious conflict is the absence of the large gametophyte clade found in all analyses of the latter study. When the monophyly of this large gametophyte clade is enforced in our analyses, we obtain a tree that is 19 steps longer is obtained. Thus, our current analyses indicate that large size and complexity of the gametophyte has evolved independently in several lineages. It should be noted that while large size and overall leaf structure are shared by these groups, there are also significant differences between them. For example, the internal structure of the stem differs in most species of *Dawsonia* (hydroids mixed with sclerenchyma) from that observed in other large gametophytes of Polytrichaceae (Smith 1971). *Dawsonia* species also have a unique type of the peristome, different from that of other peristomate Polytrichaceae. On the other hand, it should be noted that the nodes of the backbone of our optimal tree obtained have very low support values, and thus, these relationships can be easily overturned if the new additional information conflicts with the current data.

Ideally, the potential of total evidence analyses that use highly conservative genes to help in situations with conflicting hypotheses derived from morphology should be “tested” in other groups of organisms with a more completely known fossil record. Our knowledge of the bryophyte fossil record has improved in the last decades (Tomescu & al., 2018), but extinct bryophytes are still scarce and poorly known as compared to other embryophytes and, thus, few are available for inclusion in this kind of studies. In comparison, seed plants are represented today only by a fraction of their former diversity, yet recent detailed studies like the one on the conifer family Araucariaceae by Escapa & Catalano (2013) demonstrate how this kind of approach can successfully yield robust phylogenetic hypotheses.

Future studies of Polytrichopsida should be aimed at incorporating all terminals into phylogenetic analyses. This is not an unrealistic goal with a group of this size, with approximately less than 200 species (Bell & Hyvönen, 2010). Ideally future analyses should also include multiple terminals of the species that have wide geographic ranges. At the same time the aim should also be to use also the data from more variable gene regions. Difficulties in aligning these regions, particularly in the case of outgroup terminals, have been the main reason for exclusion of outgroups from previous analyses of relationships within the group (e.g. Bell & Hyvönen, 2010) that rooted trees with *Alophosia azorica* on the basis of prior studies using conserved regions and wider sampling (e.g. Bell & Hyvönen, 2008). Ideally, future analyses can be performed using direct optimization (Wheeler, 1996) in order to avoid unwarranted assumptions about homology between nucleotide positions that occur when alignment and phylogenetic analyses are separated. Of equal importance, additional fossils, some of which await detailed examination and description, need to be included in future analyses (e.g., Bippus & al., 2018). Whether these fossils will affect the overall pattern of phylogeny presented here remains to be seen. As we have seen with *Eopolytrichum*, in a group that is obviously very ancient, we cannot assume, *a priori*, that the currently known fossils, which go back in time less than 140 Ma, represent the oldest lineages of the group.

## Acknowledgements

We thank Kevin Nixon and Willi Hennig Society for making the programs Winclada and TNT freely available.

## Appendix

GenBank accessions nos. of the sequences used in the analyses supplementing those used in Bell & Hvyönen (2010). Outgroup terminals listed first, followed by the ingroup and the nos. of their *rbc*L used in the analysis.

|  | nc 18S | cp *rbcL* | *rps4* | mt *nad5* |
| --- | --- | --- | --- | --- |
| *Andreaea rupestris* | U18490 | AB469555 | AF478248 | AJ001227 |
| *Buxbaumia aphylla* | Y17603 | GQ368610 | AF231897 | KC662862 |
| *Diphyscium foliosum* | Y17765 | AF478220 | AF223034 | AY312874 |
| *Funaria hygrometrica* | X74114 | AF226818 | AJ250120 | JF501564 |
| *Oedipodium griffithianum* | AF228668 | AF478202 | AF478255 | AY312880 |
| *Sphagnum palustre* | AF126290 | AF231887 | MF362454 | KC662871 |
| *Tetraphis pellucida* | Y17604 | U87091 | AY908021 | KC662872 |
| *Timmia sibirica* | AF023678 | AJ275166 | AF023775 | -- - |

*Alophosia azorica* AY312924, *Atrichopsis compressa* EU927307, *Atrichum angustatum* DQ645986, *A. crispum* KP881775, *A. undulatum* AB917062, *Bartramiopsis lescurii* AF208409, *Dawsonia papuana* AF208410, *D. superba* AY118237, *Dendroligotrichum dendroides* AF208411, *Hebantia rigida* AY118240, *Itatiella ulei* AF208412, *Lyellia aspera* AF208413, *L. crispa* JX241626, *Notoligotrichum australe* AF208414, *N. crispulum* GU569425, *Oligotrichum hercynicum* AY118243, *O. parallelum* AF208415, *Pogonatum camusii* KU852692, *P. contortum* AY118247, *P. nudiusculum* ////////, *P. pensilvanicum* AY118253, *P. piliferum* ////////, *P. sinense* DQ120779, *P. urnigerum* AY118256, *Polytrichadelphus magellanicus* AY118257, *Polytrichastrum alpinum* GU569464, *P. lyallii* AY118241, *Polytrichum formosum* AY118259, *P. longisetum* AY118260, *P. commune* U87087, *P. piliferum* AY118263, *Psilopilum laevigatum* AF208416, *Steereobryon subulirostrum* AY118265

